# Collaborative workflow between pathologists and deep learning for evaluation of tumor cellularity in lung adenocarcinoma

**DOI:** 10.1101/2022.01.11.475587

**Authors:** Taro Sakamoto, Tomoi Furukawa, Hoa H.N. Pham, Kishio Kuroda, Kazuhiro Tabata, Yukio Kashima, Ethan N. Okoshi, Shimpei Morimoto, Andrey Bychkov, Junya Fukuoka

**Affiliations:** Department of Pathology, Nagasaki University Graduate School of Biomedical Sciences, Nagasaki, Japan; Department of Pathology, Kameda Medical Center, Kamogawa, Japan; Department of Pathology, Awaji Medical Center, Sumoto, Japan; Innovation Platform and Office for Precision Medicine (iPOP), Graduate School of Biomedical Sciences, Nagasaki University, Nagasaki, Japan

**Keywords:** artificial intelligence, digital pathology, deep learning, lung adenocarcinoma, tumor cellularity

## Abstract

Owing to the high demand for molecular testing, the reporting of tumor cellularity in cancer samples has become a mandatory task for pathologists. However, the pathological estimation of tumor cellularity is often inaccurate.

We developed a collaborative workflow between pathologists and artificial intelligence (AI) models to evaluate tumor cellularity in lung cancer samples and prospectively applied it to routine practice. We also developed a quantitative model that we validated and tested on retrospectively analyzed cases and ran the model prospectively in a collaborative workflow where pathologists could access the AI results and apply adjustments (Adjusted-Score). The Adjusted-Scores were validated by comparing them with the ground truth established by manual annotation of hematoxylin-eosin slides with reference to immunostains with thyroid transcription factor-1 and napsin A. For training, validation, retrospective testing, and prospective application of the model, we used 40, 10, 50, and 151 whole slide images, respectively.

The sensitivity and specificity of tumor segmentation were 97% and 87%, and the accuracy of nuclei recognition was 99%. Pathologists altered the initial scores in 87% of the cases after referring to the AI results and found that the scores became more precise after collaborating with AI. For validation of Adjusted-Score, we found the Adjusted-Score was significantly closer to the ground truth than non-AI-aided estimates (p<0.05). Thus, an AI-based model was successfully implemented into the routine practice of pathological investigations. The proposed model for tumor cell counting efficiently supported the pathologists to improve the prediction of tumor cellularity for genetic tests.

## 1. Introduction

The rapid growth of artificial intelligence (AI) in recent years, especially deep learning (DL), has provided significant advancements in numerous fields including medicine. Among several machine learning methods, the convolutional neural network (CNN) has emerged as the most suitable method for medical image analysis^1-4^. Trials using CNNs to assist physicians with diagnosis, treatment, or even prognosis have produced extremely promising results^5-9^ that represent a veritable explosion in the field of digital medicine.

In pathology, CNNs have been used to analyze various tissues and detect tumor regions to support histopathological diagnosis^10-15^. Several studies have demonstrated that CNNs can provide judgments equivalent to those of pathologists, or even exceed them in certain special tasks^12,14^ However, the implementation of AI in pathology still faces challenges such as the high rate of false positives and false negatives, and the training procedure of the algorithms for optimizing the balance between these two kinds of results^16^. Thus, collaboration between the pathologists and DL may be the most suitable approach to overcome these obstacles as we still do not completely understand the nature of the “black box” inside the DL training process^17^. The implementation of DL in clinical workflow including evidence of its safety for patient healthcare is an important issue that needs to be addressed^18,19^.

For decades, lung cancer has had the lowest survival rate among all types of cancers in humans^20^. Recent years have experienced revolutionary developments in cancer treatment, such as “personalized” molecular therapies and the introduction of checkpoint inhibitors, with several studies presenting encouraging evidence to their efficacy^21,22^. The increased prevalence of molecular testing, e.g., evaluation of the mutation status of *EGFR* and *RAS*, has yielded the percentage of tumor cells in the test sample being identified as a statistic of prime importance for ensuring the success of the test^23-25^. The minimum required percentage of tumor cells in a sample is dependent on the analytical sensitivity of the platform conducting the tests and considerably varies between the platforms^25,26^. Thus, the evaluation of tumor cellularity by pathologists, i.e., the percentage of tumor cells in the sample, is considered critical. However, recent studies have reported that a high variability and low reproducibility exist across individual pathologists^27-30^. Precise tumor cellularity is required to glean essential information from molecular testing, which can assist pathologists to evaluate whether the mutations arise entirely from the tumor cells, from a minor component of tumor cells, or nearby “benign-looking” cells. The development of an effective and objective method is imminently required to precisely estimate tumor cellularity.

Several studies have been conducted to detect, discriminate subtypes of, and predict the mutations of lung cancers using histological features of tumor cells^9,18,31-36^. However, to the best of our knowledge, no study has focused on measuring tumor cellularity using DL—which is expected to aid pathologists in accurately determining the tumor cellularity value for molecular testing. In this report, we describe our experience with AI implementation in a clinical lung cancer workflow, in which a DL algorithm was developed and applied both retrospectively and prospectively to assess its impact on the quality of molecular testing.

## 2. Materials and Methods

### 2.1. Study cohorts

The current study protocol was approved by the Institutional Review Board of Nagasaki University Hospital (#190218282). We designed the study in three phases: an algorithm development phase, a retrospective phase, and a prospective phase (Figure 1). In the algorithm development phase, a CNN model was constructed and validated to measure tumor cellularity. In the retrospective phase, the model was retrospectively tested on cases, and in the prospective phase, its efficacy was evaluated in prospective cases. For the algorithm development and retrospective phases, a total of 100 hematoxylin and eosin (H&E)-stained transbronchial lung biopsy (TBLB) slides (one slide per case) diagnosed with lung adenocarcinoma were collected from Nagasaki University Hospital, Nagasaki, Japan. Of these, 50 whole slide images (WSIs) from 2014 to 2017 were used as a dataset for model development, and 50 slides from 2017 to 2018 were used in the retrospective testing study (retrospective phase). For the prospective phase, 151 slides from 2019 to 2020 were used for the review. These 151 slides comprised five distinct types of sampling methods: transbronchial biopsy/transbronchial lung biopsy (TBB/TBLB), core needle biopsy (CNB), surgical resection, transbronchial needle aspiration/transbronchial aspiration cytology (TBNA/TBAC), and cell block. These slides were scanned using an Aperio Scanscope CS2 digital slide scanner (Leica Biosystems, Buffalo Grove, IL) with a 40x objective lens (0.25 µm/pixel) for the retrospective phase. Similarly, 151 slides for the prospective phase were scanned using an Ultra Fast Scanner (Philips, Amsterdam) with a 40x objective lens (0.25 µm/pixel). Thereafter, digital slides were imported into HALO version 2.2 (Indica Labs, Corrales, CA), which included HALO AI (CNN VGG network) and the HALO Image Analysis program.

**Figure 1.**
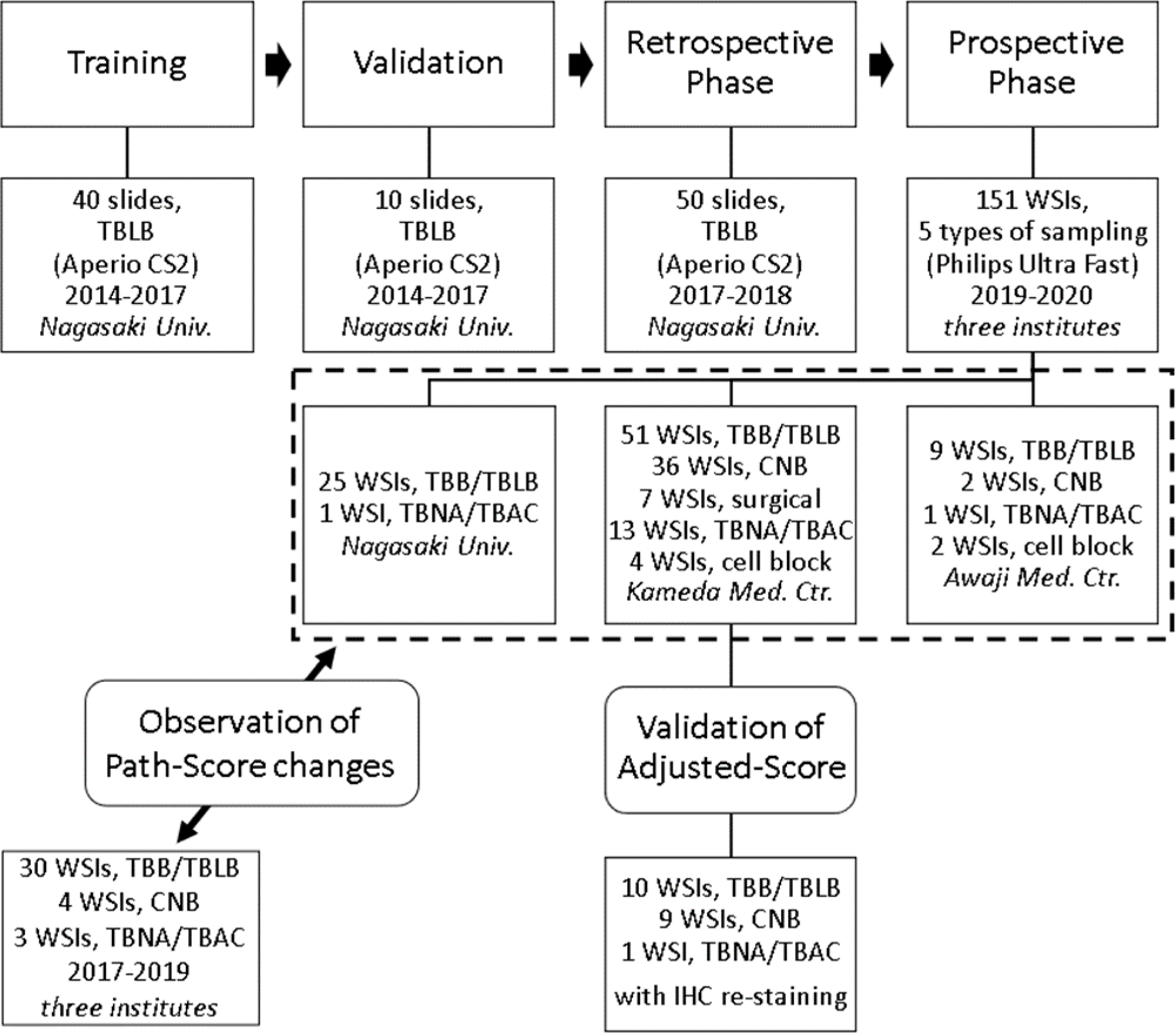
Flowchart of study dataset. The study dataset was designed in three phases: algorithm development phase (training and validation), retrospective phase, and prospective phase. This flowchart includes the number of each dataset, types of sampling, name of scanners, year, and the institution’s name. WSI: whole slide image; TBB/TBLB: transbronchial biopsy/transbronchial lung biopsy; CNB: core needle biopsy; Surgical: surgical resection; TBNA/TBAC: transbronchial needle aspiration/transbronchial aspiration cytology; Nagasaki Univ.: Nagasaki University Hospital; Kameda Med. Ctr.: Kameda Medical Center; Awaji Med. Ctr.: Awaji Medical Center; IHC: immunohistochemical.

### 2.2. Algorithm Development Phase: Model development and verification

We enrolled 50 TBLB slides as a dataset for DL model development. Among the 50 cases, 40 were used for training, and the remaining 10 cases were used for validation. The tumor region and nontumor region annotations for the training data were comprehensively identified at the region level by a trainee and reviewed by two expert pulmonary pathologists (J.F. and K.T.). In total, 10,644 annotations (5625 tumor regions, 5019 nontumor regions) were made. As the resolution was set at 0.2517 µm/pixel, 0.01 mm^2^ patches were randomly and automatically generated, resulting in 3,085,569 patches, to be used as a training dataset for the DL model.

Subsequently, the model was validated on the 10 cases of the validation dataset. The annotations were made in the same manner as those for the slides, which were used as the ground truth for comparison with the predictions of the DL model. All the annotations were made based on the consensus of three pathologists (J.F., K.K., and K.T.), and additional immunohistochemical staining of thyroid transcription factor-1 (TTF-1), napsin A, p40, and CK14 was employed to highlight cell populations. The 10 validation case images were divided into 0.01 mm^2^ patches, producing 14,611 clipped patches. Each patch was evaluated by overlaying the annotations made by the pathologists on those generated by the DL model, after which the areas of true positive, true negative, false positive, and false negative were evaluated. Based on this overlay data, the sensitivity, specificity, and accuracy of the tumor-region detection by the model were calculated.

We used HALO Image Analysis to detect all nuclei in the tissue for measuring the tumor cellularity (i.e., tumor cell percentage). The parameters for nuclei detection were selected by executing a nuclei detection algorithm on 10 randomly selected cases from the training dataset, including 5 patches of both tumor and nontumor regions. We compared the nuclear counting performance obtained from HALO with the manually conducted counts and selected the parameters with the highest accuracy in comparison to the manually counted results.

### 2.3. Retrospective Phase: Testing study of DL model in retrospective cases

An alternative set of 50 TBLB WSIs was used to retrospectively test the model by comparing pathologist- and AI-generated predictions. First, a representative fragment of each case was selected. Four pathologists, J.F., A.B., Y.K., and H.P., manually estimated the tumor cellularity by reviewing the virtual slides. The tumor regions determined by the DL model constructed in the algorithm development phase were subsequently combined with the data obtained from the HALO Image Analysis software to calculate the percentage of nuclei detected within the highlighted tumor areas, thus evaluating the overall tumor cellularity (Figure 2).

**Figure 2.**
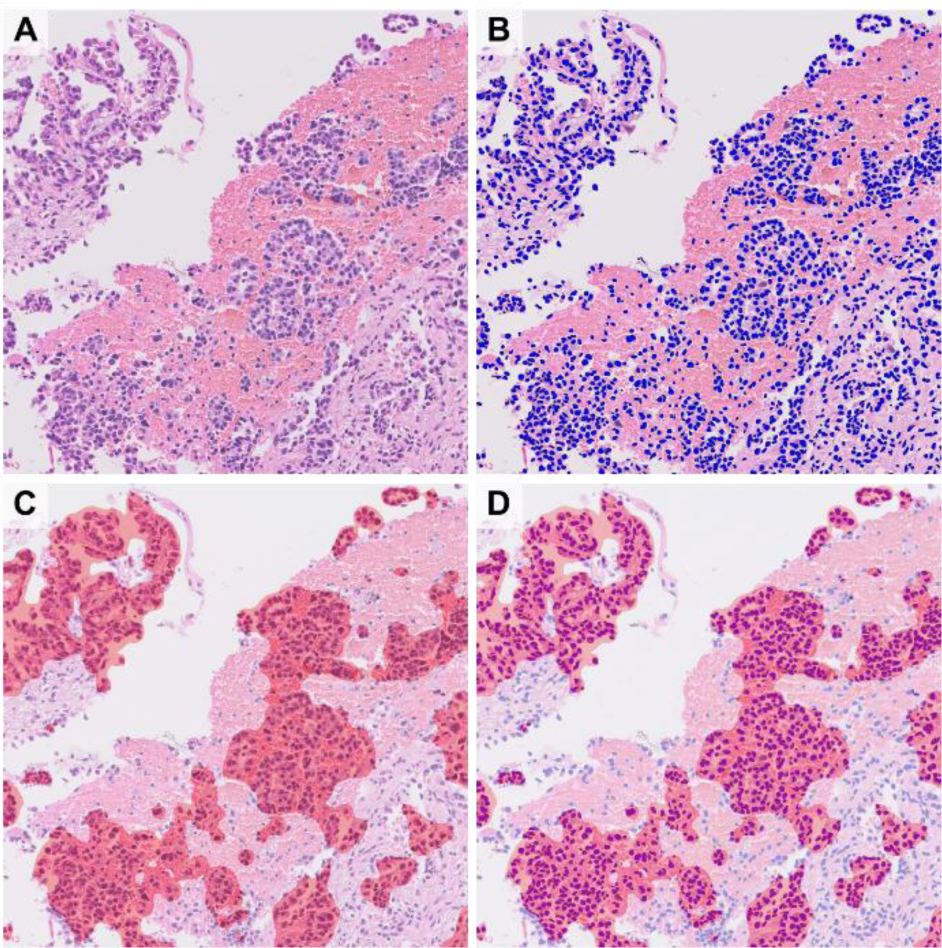
Calculation of tumor cellularity using two algorithms. Original H&E image (a) was analyzed using two distinct algorithms. Nuclei on H&E image were masked as blue markers (b), and tumor clusters were segmented as red masking (c). Combining (b) with (c), the total number of nuclei (number of blue markers) and the number of tumor nuclei (number of blue markers within red mask) could be obtained (d), so we can calculate the tumor cellularity of samples.

Based on the perceived level of accuracy by a group of pathologists, these DL model-generated tumor cellularity estimates were sorted into three groups: good (≥ 90%), fair (70– 90%), and poor (< 70%) following the consensus of attending pathologists. Thereafter, the ground truth was established using a combination of the pathologists’ annotations for tumor regions and the HALO Image Analysis detection of nuclei for tumor cellularity. A statistical comparison between the results of the DL model and pathologists was performed based on the deviation (absolute value of difference) between these results and the ground truth.

### 2.4. Architecture of VGG-16 model

Upon analyzing the tissue using the developed DL model and detecting the tumor regions, the HALO Image Analysis detected the nuclei within the labeled tumor region as well as in the entire specimen. It used the Caffe engine and a fully convolutional version of the VGG architecture with all padding removed. The training was conducted on 435 × 435 patches at the defined resolution, which were generated by selecting a random class (with equal probability for each class), a random image containing annotations for the selected class (with equal probability), and a random point inside a region of the selected class and image. The patches were cropped surrounding the selected point and were further augmented with random rotations and shifts in hue, saturation, contrast, and brightness. The model was pre-trained on ImageNet, and thereafter, trained for the defined number of iterations using RMSProp (delta of 0.9) with a learning rate of 1e-3, which was reduced by 10% every 10,000 iterations along with an L2 regularization of 5e-4. As there was no padding in the model during analysis, the tile size was increased to 1867 × 1867 without altering the output or increasing the performance. NVIDIA GTX TITAN X graphics card was used as the GPU required for HALO AI.

### 2.5. Prospective Phase: Routine application of DL model in prospective cases

#### 2.5.1. Creation of workflow

From April 2019 to September 2020, biopsies and surgical resections of pulmonary adenocarcinoma cases from three institutes (Nagasaki University Hospital, Kameda Medical Center, and Awaji Medical Center) within our network^18^ were prospectively enrolled in this study. All the institutions are fully digitized, i.e., glass slides were scanned before assigning the cases to the pathologist. The designed workflow is presented in Figure 3. Initially, the WSIs of the suspected adenocarcinoma cases were immediately examined by the assigned pathologists, and the WSIs containing the highest number of tumor cells were selected as per the pathologists’ assessment. These selected images were downloaded by a member of the analysis team and converted to the pyramid TIFF format following anonymization. For certain cases, the images were cropped. The tumor cells were annotated to enclose the region of interest (ROI) (Figure 3). The WSIs were evaluated using the trained algorithm, and the results were shared at the sign-out sessions of the following day. At the sign-out sessions, the pathologists were blinded to the results of the AI analysis and were asked to estimate the tumor cellularity of the specimens; their answers were averaged to produce the Path-Score. Subsequently, the pathologists visually reviewed the results of the AI analysis represented by the automated tumor detection map and nuclear detection overlay (AI-Score). The pathologists determined the final tumor cellularity (Adjusted-Score) by adjusting the AI-Score as they deemed fit. As described earlier, the quality of AI analysis was categorized into three levels; the representative images for each level are presented in the supplementary data (Supplementary figure 1).

**Figure 3.**
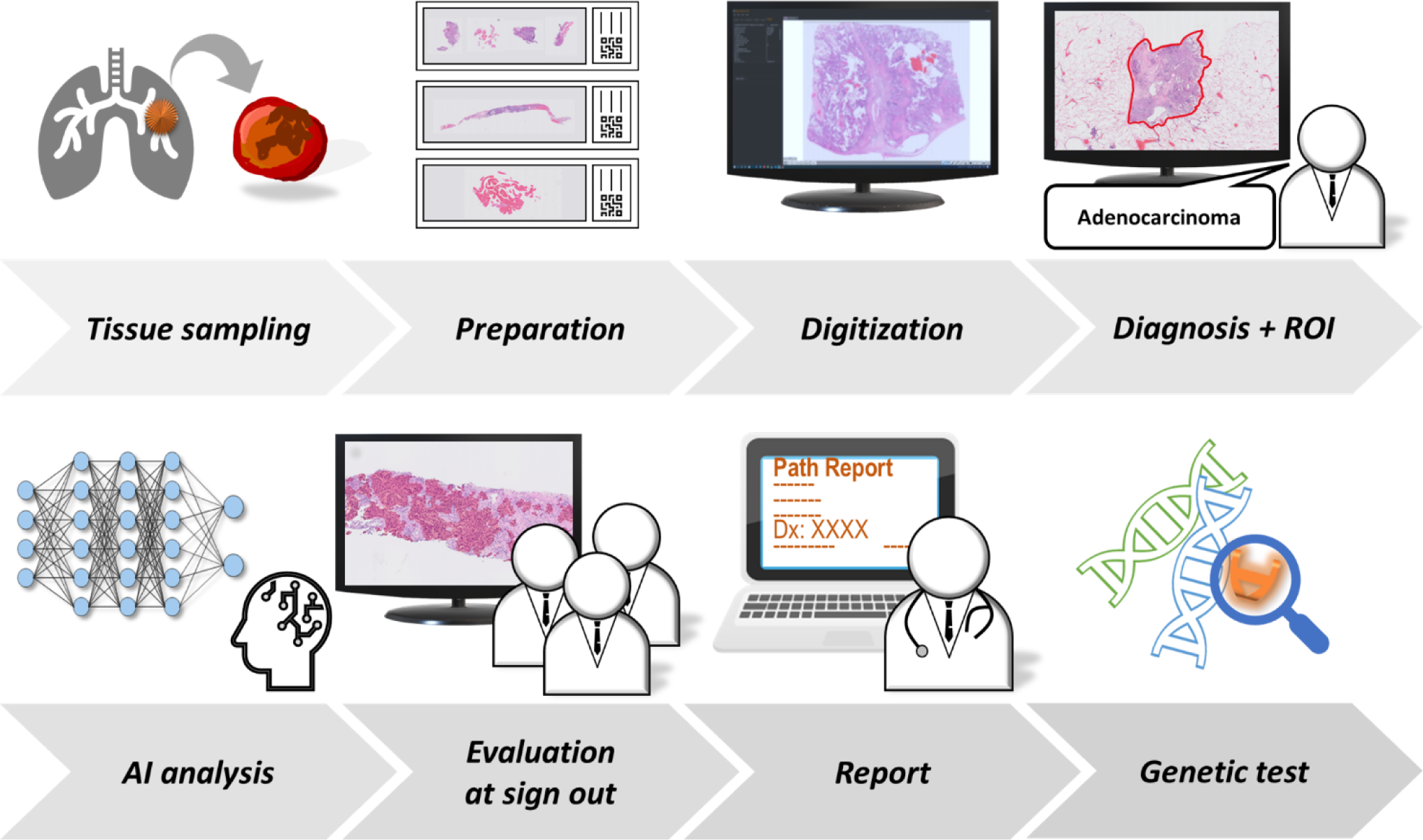
Synergistic workflow between pathologists and AI model. The proposed approach significantly improved the pathologists’ workflow by enabling them to diagnose using digital images. The approach comprised routine operation of AI model and evaluation of AI analysis results by pathologists. Starting with tissue sampling, specimen preparation, and digitization, the cases diagnosed as adenocarcinoma by pathologists were assigned with ROIs as necessary and subjected to analysis of tumor cellularity by AI. After the analysis, the cases were evaluated and modified by pathologists in a sign-out session and Adjusted-Scores were recorded in path report and forwarded to genetic testing lab.

#### 2.5.2. Adjustment to numbers indicated by AI (Adjusted-Score)

The cases requiring further major or minor adjustment of the AI-Score were mathematically processed. For instance, in Supplementary figure 2, the recognition of tumor cells by AI was considered as 30% less than the actual value. Therefore, based on the consensus of the pathologists, we added 30% to the AI-Score to obtain the Adjusted-Score.

#### 2.5.3. Validation of Adjusted-Score using immunohistochemical staining

Twenty adjusted cases were randomly selected, and the tumor cells were manually annotated. The original H&E slides were de-stained with hydrochloric acid/ethanol solution and immunohistochemically re-stained with a cocktail of antibodies for TTF-1 and napsin A (ADC cocktail, Pathology Institute Corp., Toyama, Japan)^37^ which are widely used markers for pulmonary adenocarcinoma. The stained slides were further scanned using the Ultra Fast Scanner. Annotations for individual tumor cells were applied based on simultaneous observations of the immunostained slides and the original H&E images (Supplementary figure 3). All the annotation data were verified by the expert pulmonary pathologist (J.F.). Ultimately, the cell count algorithm was applied to the annotated area, and the tumor cellularity values from those twenty cases were used as the ground truth. These numbers were compared to the original Path-Scores and Adjusted-Scores.

### 2.6. Statistical analysis

The chronological variation in the scores assigned by the individual pathologist through the prospective phase was examined using logistic regression analysis. Moreover, we examined whether the tendency of overestimation by more than 20% was ameliorated after observing 20 cases. We also employed the generalized Wilcoxon test to compare variations in the individual pathologist’s tendency to overestimate scores between retrospective and prospective analysis. A Wilcoxon signed-rank test was conducted to assess the adjustments to the AI-Score (Adjusted-Score) in 20 out of 151 cases in the prospective phase, wherein a two-sided *p-value* < 0.05 was considered statistically significant.

## 3. Results

### 3.1. Algorithm Development and Retrospective Phase

To validate the proposed DL software in the algorithm development phase, a model with a cross entropy of 0.16 was adopted as the test model. The validation set contained 10 whole slide images that were divided into 14,611 patches of 0.01 mm^2^ to match the size of the training patches. Among these 14,611 patches, 7771 were classified by the DL model as positive for the tumor regions, and 7630 regions corresponded with the pathologists’ positive annotation, i.e., true positives. Conversely, 6024 out of 6840 patches were true negatives that were classified as negative by the DL model as well. The overall sensitivity, specificity, and accuracy were 97.1%, 87.0%, and 93.5%, respectively. To evaluate the nuclear recognition, the HALO Image Analysis results were compared with the pathologists’ exact manual count in 10 ROIs, which revealed an accuracy of 98.5%.

In evaluating tumor cellularity in 50 randomly selected cases, the mean deviation from the ground truth among the four participating pathologists was 15%, whereas the mean deviation of the results obtained by the proposed DL model from ground truth was 6%. Among these 50 cases, 29 were categorized as good, 12 were categorized as fair, and 9 were categorized as poor. In 19 of the 29 cases categorized as “good,” the DL model outperformed all the participating pathologists (Figure 4, Supplementary table 1). The mean deviation of the DL model in these “good” cases was 3%, whereas the mean deviation for the pathologist estimations was 16%. For the 12 “fair” cases, the DL model deviated 4% on average from the ground truth, whereas the pathologists differed from the ground truth by 14% on average. For the 9 “poor” cases, the mean deviations were 15% and 14% for the DL model and pathologists, respectively. The results show that the consensus judgment of pathologists generally deviated from ground truth by approximately 15%.

**Figure 4.**
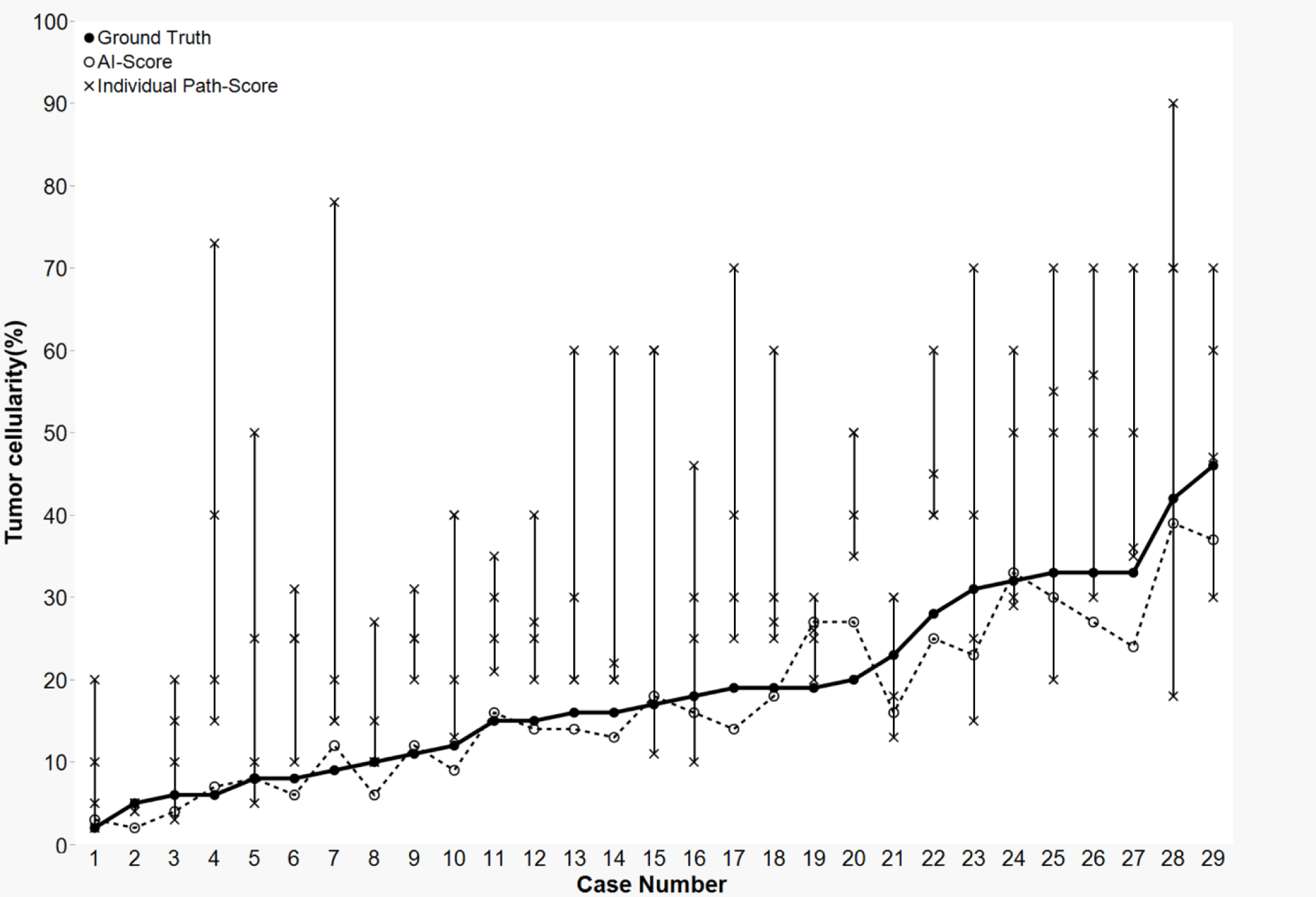
Inter-observer variability and inconsistency of pathologist’s tumor cellularity estimates in 29 cases assessed as “good” in retrospective phase. A line plot displaying Ground Truth (solid line), AI-Score (dotted line), and individual Path-Scores (vertical line). In 19 out of 29 cases, AI-Score outperformed all participating pathologists’ assigned Path-Scores.

### 3.2. False positives and false negatives

False positives mainly arise from erroneous detection of bronchial epithelium, macrophages, lymphocytic aggregates, bronchial cartilages, and anthracotic pigments as tumor cells (Supplementary figure 4). In certain instances, the proposed segmentation model could not appropriately identify mildly atypical cells and low nuclear-cytoplasmic ratio lesions such as lepidic growths or invasive mucinous adenocarcinomas (Supplementary figure 4). Such false negatives were observed in the lesion, wherein the tumor cells were spindle-shaped and had infiltrated the stroma in necrotizing carcinoma tissue areas or in out-of-focus areas missed by the scanner.

### 3.3. Assessing level of classification in Prospective Phase

A total of 151 samples were prospectively analyzed in the prospective phase of this study. Among these samples, 26 were acquired from Nagasaki University Hospital, 111 from Kameda General Hospital, and 14 from Awaji Medical Center (Figure 1). Following the consensus of the participating pathologists, the AI segmentation of the samples was labeled by pathologists as follows: 80 good (53%), 38 fair (25%), and 33 poor (22%). The pre-test data collected in the retrospective analysis during the initiation of the routine usage of AI displayed a tendency for overestimation (i.e., ≥ 1% increments) by an individual pathologist (overestimation in 21/37). The prospective data displayed a similar frequency of overestimation and underestimation: overestimation in 67/151 cases (44.3%) and underestimation in 65/151 cases (43%), as depicted in Supplementary figure 5. Based on the generalized Wilcoxon test, the pathologist tended to significantly overestimate the tumor cellularity to a greater extent in the 37 retrospective pre-test samples than the 151 prospective samples (*p* < 0.005).

The pathologist who led this study was involved in the sign-out of all the 151 cases enrolled herein. Before reaching 20 cases, the pathologist tended to overestimate the tumor cellularity by more than 20%, resulting in a large deviation between the individual Path-Score and Adjusted-Score. However, once the pathologist was able to refer to the AI segmentation data from the 20 cases, the pathologist realized that he tended to overestimate (Figure 5). Based on logistic regression analysis, the pathologist’s score did not tend to deviate from the Adjusted-Score by more than ±20% onward case no. 20 (*p* = 0.019). This pathologist’s Path-Score deviated from the Adjusted-Score in 132 out of 151 samples, which signified that the AI segmentation data caused the pathologist to reconsider their first estimate of the tumor cellularity in 87% of the samples.

**Figure 5.**
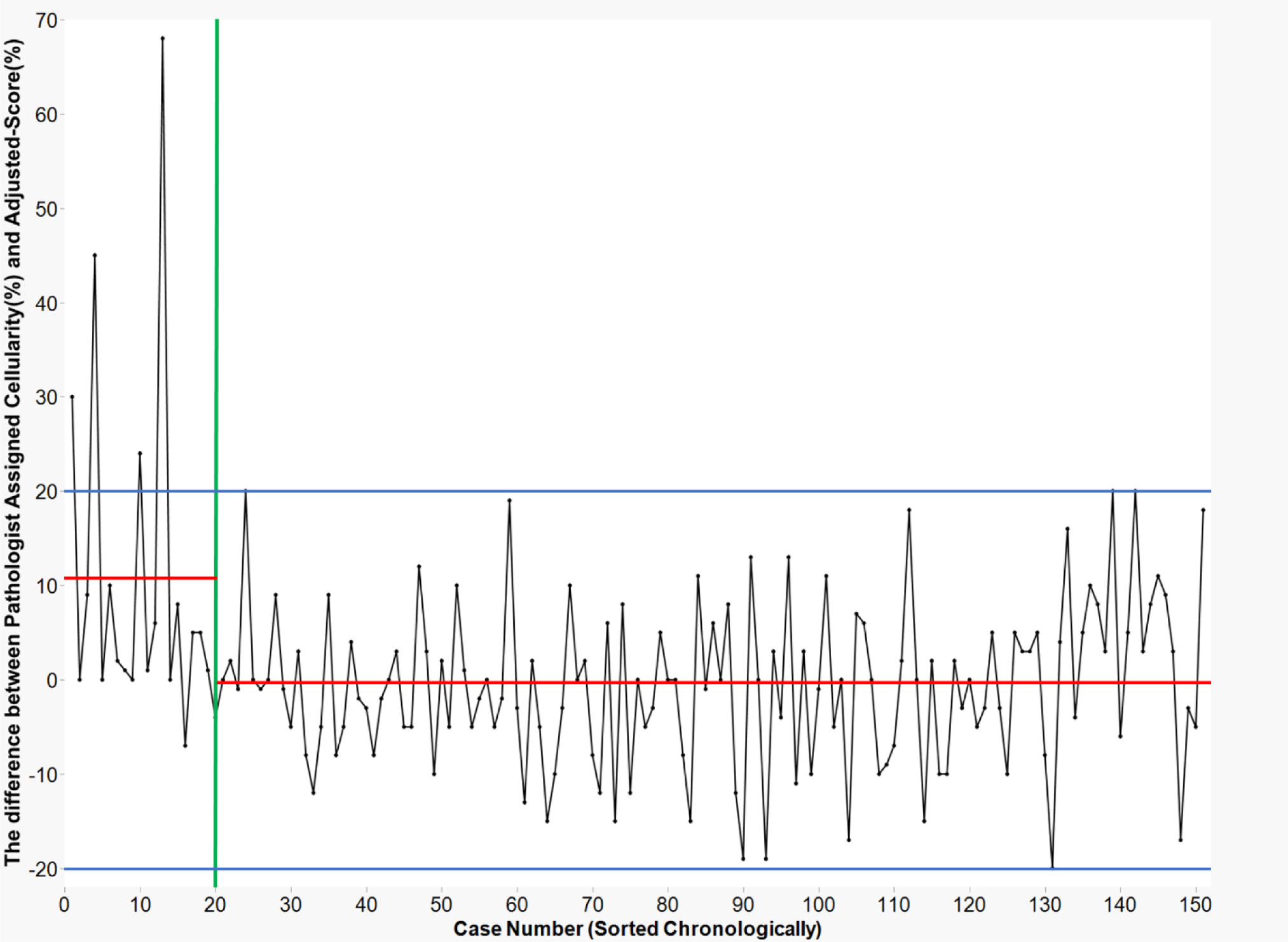
Improvement of cellularity estimation. Chronological variation of deviation between Adjusted-Score and Path-Score for an individual pathologist. In the early stage, overestimation exceeding 20% deviation from Adjusted-Score appeared in certain cases, but the Path-Scores stabilized. Although the pathologists supervised the AI, they could likewise learn from it. These data not only implied that the human-in-the loop workflow effectively improved the pathologist’s assessment, but also highlighted the requirement of AI aid (despite increased trials of cellularity estimates, the pathologist’s score varied). Red lines: average difference between individually-assigned cellularity and Adjusted-Score. Separate averages are shown for case numbers 1 to 19 (average: 10.95), and from 20 on (average: -0.598). Blue lines: ±20% is shown as a cut-off range in percent difference. Green line: case number 20.

### 3.4. Comparison with ground truth

The total number of individual tumor cluster level annotations in the 20 randomly selected ground truth cases was 4527. This ground truth was established by manually counting the tumor nuclei on H&E stained slides while referencing the same slide re-stained with TTF-1/napsin A. The average number of annotations was 226.35 per sample (range, 33– 526). The mean deviation between the ground truth and Adjusted-Score was 3.08%. The Adjusted-Score was within 5% of the ground truth in 80% of the test cases (16/20), and within 10% in all the cases. The mean deviation of the pathologist’s Path-Score to ground truth was 7.12%, confirming that the Adjusted-Score was generally closer to the ground truth (*p* = 0.009, Wilcoxon signed-rank test) (Figure 6, Supplementary figure 6). The Adjusted-Score was also superior to the scores obtained by the consensus of pathologists (*p* = 0.032, Wilcoxon signed-rank test). Among the 20 test cases, the AI segmentation data were categorized as “good” in 4 cases, “fair” in 12 cases, and “poor” in 4 cases. The median deviation from the ground truth in each level was 3.2%, 2.1%, and 4.0%, respectively (Supplementary figure 7).

**Figure 6.**
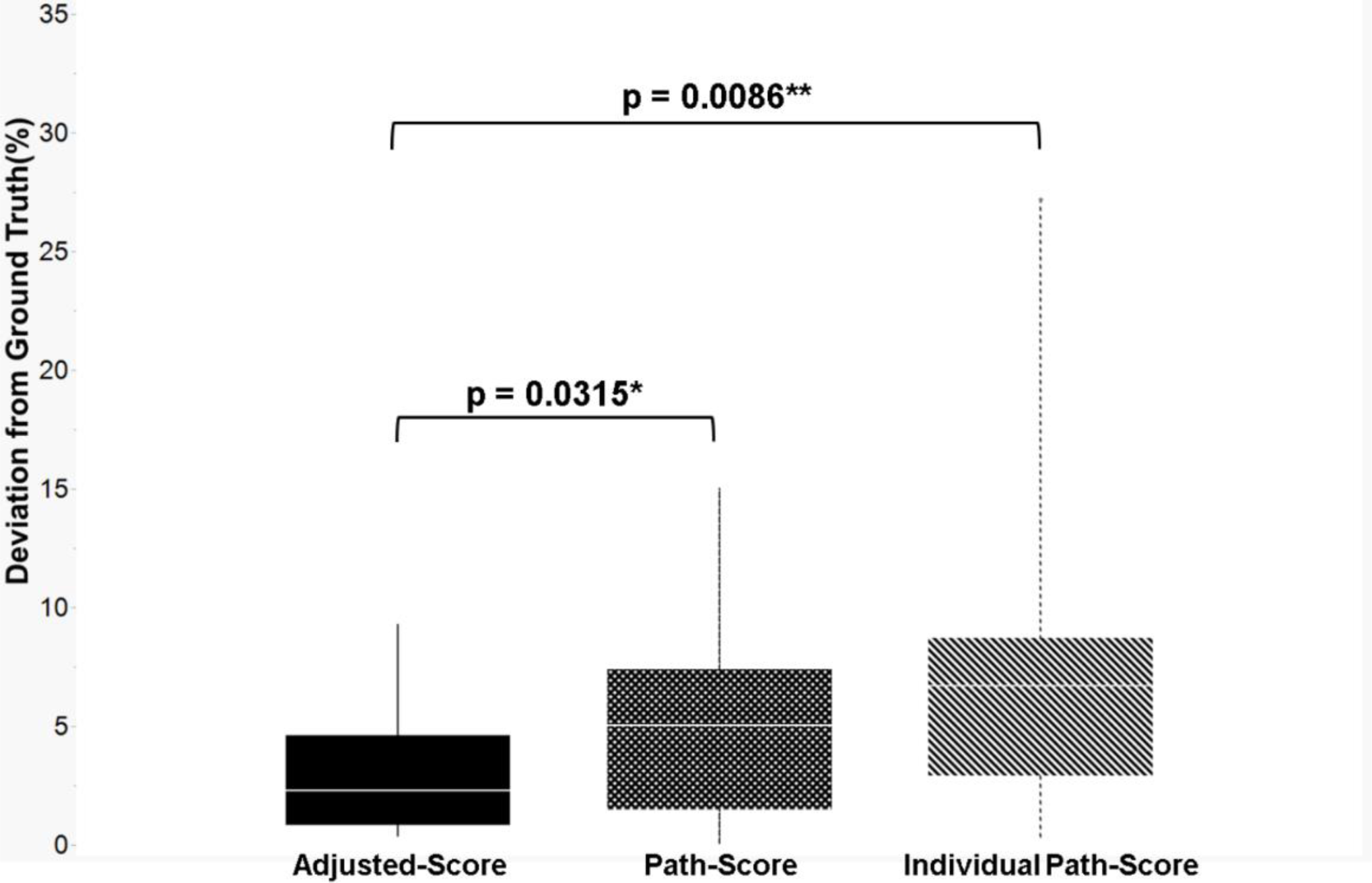
Deviation from ground truth. The box plot displaying Adjusted-Score was significantly closer to the ground truth than both the Path-Score and Individual Path-Score for a pathologist. The proposed AI-based tumor cellularity adjustment attained less than 10% deviation in all the cases, and less than 5% deviation in three quarters of the cases.

### 3.5. Assessment of level of classification per sampling modality and among institutions

The proportion of samples categorized by the model as “good” for each sampling method was 53% for TBB/TBLB, 39% for CNB, 57% for surgical, 93% for TBNA/TBAC, and 33% for cell block (Table 1). The proposed model performed especially well with the TBNA/TBAC samples. The mean adjustment from AI-Score to Adjusted-Score for each modality was –19.33% for the cell block, +2.53% for CNB, –0.29% for surgical, +1.41% for TBB/TBLB, and +2.33% for TBNA/TBAC (Supplementary table 2). The false positives were highly prominent in the cell block samples (standard deviation of 31%), and the individual cells contained in the cell block samples could not be easily identified, especially mesothelial cells and macrophages floating in the pleural fluid in the thoracic cavity.

**Table 1.** Classification level per sampling method. TBB/TBLB: transbronchial biopsy/transbronchial lung biopsy; CNB: core needle biopsy; Surgical: surgical resection; TBNA/TBAC: transbronchial needle aspiration/transbronchial aspiration cytology

| Sampling | n | Good | Fair | Poor |
| --- | --- | --- | --- | --- |
| TBB/TBLB | 85 | 52.9% (45) | 24.7% (21) | 22.4% (19) |
| CNB | 38 | 39.5% (15) | 34.2% (13) | 26.3% (10) |
| Surgical | 7 | 57.1% (4) | 42.9% (3) | – |
| TBNA/TBAC | 15 | 93.3% (14) | 6.7% (1) | – |
| Cell Block | 6 | 33.3% (2) | – | 66.7% (4) |
| Total | 151 | 53% (80) | 25% (38) | 22% (33) |

The percentage of samples rated as “good” from each institution were as follows: 73% from Nagasaki University Hospital (19/26), 48% from Kameda Medical Center (53/111), and 57% from Awaji Medical Center (8/14) (Supplementary table 3). Upon including the samples rated as “fair,” the percentages from each institution were 88% from Nagasaki University Hospital (23/26), 76% from Kameda Medical Center (84/111), and 79% from Awaji Medical Center (11/14) (Supplementary table 3).

## 4. Discussion

We developed a DL image analysis model that yielded tumor cellularity values by integrating a tumor region segmentation model and a nuclear counting algorithm. In addition, we developed a prospective trial workflow and conducted a prospective study to compare the pathologists’ estimates before and after referring to the AI segmentation data.

In our study, pathologists were able to refer to the AI-Scores and adjust them to provide more accurate estimations than the original pathologists’ consensus estimates (Path-Score). In particular, 87% of the 151 test samples involved a pathologist altering the earlier estimation after referring to AI data. Thus, this is a concrete example of incorporating AI in daily practice to provide a more accurate pathological diagnosis. We demonstrated that AI-aided diagnosis improved the pathologists’ judgment. This primarily materialized in the form of improving large overestimations in the early stages and increased the proportion of underestimations, which did not exhibit significant improvement afterwards, regardless of the enhanced case experience (Figure 5). As the human judgment is limited in this context, this may be a rationale for recommending AI-aided diagnosis. It proved that even for suboptimal accuracy of the AI, the pathologists could visually assess the results of the segmentation model and adjust the calculation of the tumor percentage to obtain a more accurate prediction of the ground truth. Upon referring to the AI data, the human could extract more accurate data, indicating that the collaboration was significant. AI may or may not be able to accurately recognize the cancer cells, and in certain instances, it did not function appropriately in the analysis of diverse images with a variety of histological reactions which is a common occurrence in pathological investigations. Decisions are ultimately taken by the pathologists, but this collaboration between the AI and pathologists—alternatively expressed as a type of human-in-the-loop^17,38,39^—is a promising direction for the future of pathological diagnosis and related tasks.

A large inter-observer variability was present in the estimates between the pathologists (Figure 4, Supplementary table 1), similar to that reported in several prior studies^27-30^ which challenged consensus-building. In this study, we adopted the average scores marked by the pathologists as the consensus. Although this averaged score was an improvement over individual pathologists’ scores, the collaborative method between the physician and the AI was ultimately the most accurate method. In clinical applications of AI, pathologists must be able to interpret the results and provide input. We overlaid the segmentation data on the WSI, reviewed the tumor cellularity percentages, verified the level of nuclear recognition, and modified the AI-Scores. Pathologists could enhance their practice by utilizing the AI model. Moreover, differences in staining and specimen preparation techniques between laboratories are some of the major barriers to the adoption of AI^40,41^. This was confirmed as the model yielded the best results on samples acquired from the institution at which training was conducted (Supplementary table 3), as well as a higher frequency of errors than on samples from other institutions. However, the input from a pathologist was able to compensate for these minor errors. We examined whether the accuracy of the Adjusted-Score varied with the decreasing classification level, but no clear deterioration was observed (Supplementary figure 7). This indicated that the specimen preparation and staining procedures conducted at various institutions did not significantly impact the tumor nuclei count obtained using the proposed model, and the AI model can be used at any institution.

There are certain limitations in this study. Although the annotations of the training data were highly accurate at the regional level, the number of cases was small. Second, in the prospective phase, the WSI scanner was altered, but the AI model was not adjusted to match the scanner.

## 5. Conclusion

We developed an AI model and a human-in-the-loop workflow to evaluate tumor cellularity. This study demonstrated that the proposed model could more accurately determine tumor cellularity than pathologists’ consensus, and additionally, implied that the pathologists can learn from the AI implementation. The collaboration between the AI and pathologists can result in a synergistic, positive feedback loop in which each side improves the other.

