## Supplementary file for "Collaborative workflow between pathologists and deep learning for evaluation of tumor cellularity in lung adenocarcinoma"

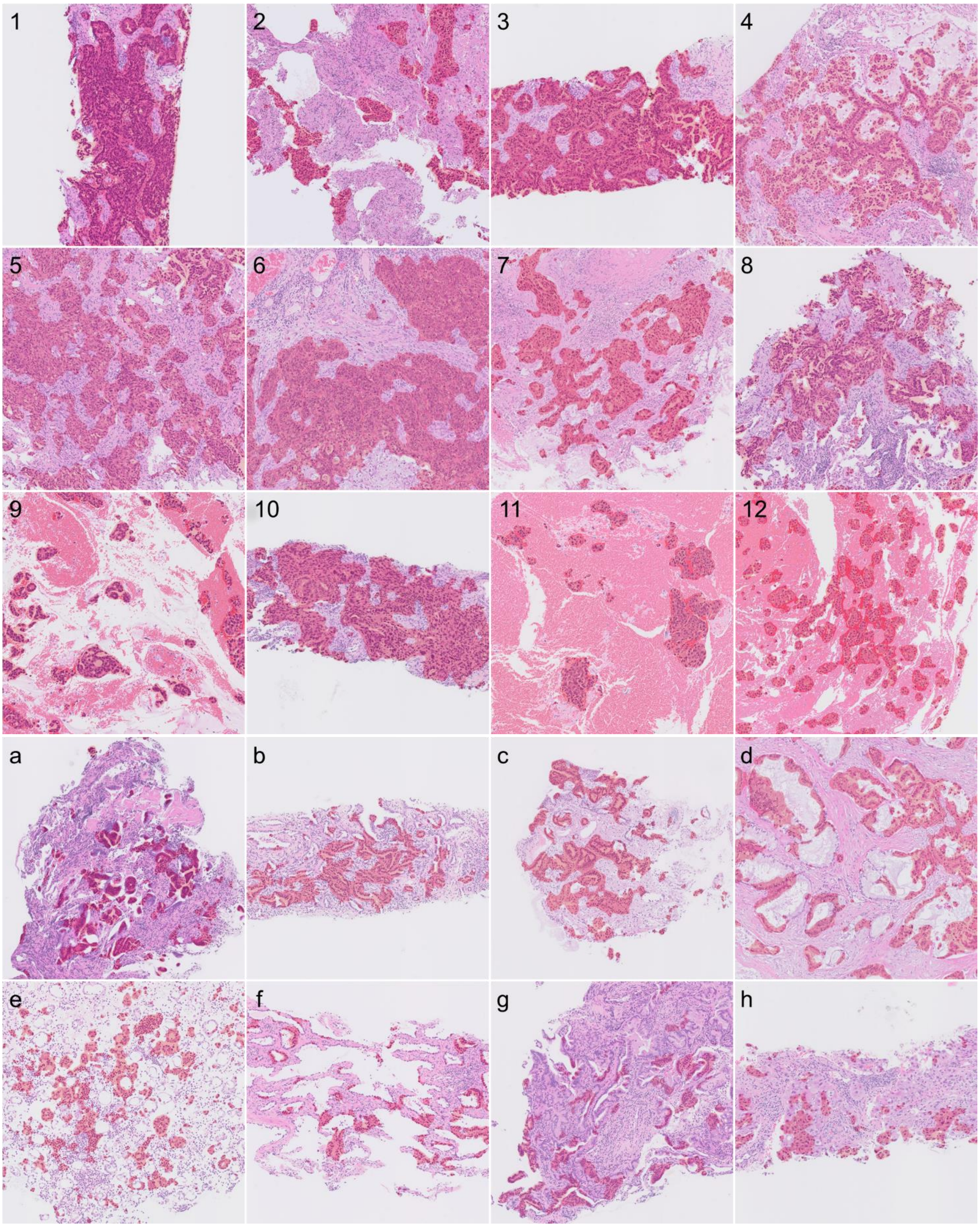


**Supplementary figure 1.** Representative images with red classification mapping judged as good, fair, and poor by pathologists.

The segmentation level of each of the cases was judged as one of the three levels: good (1–12), fair (a–d), and poor (e–h). (40x)

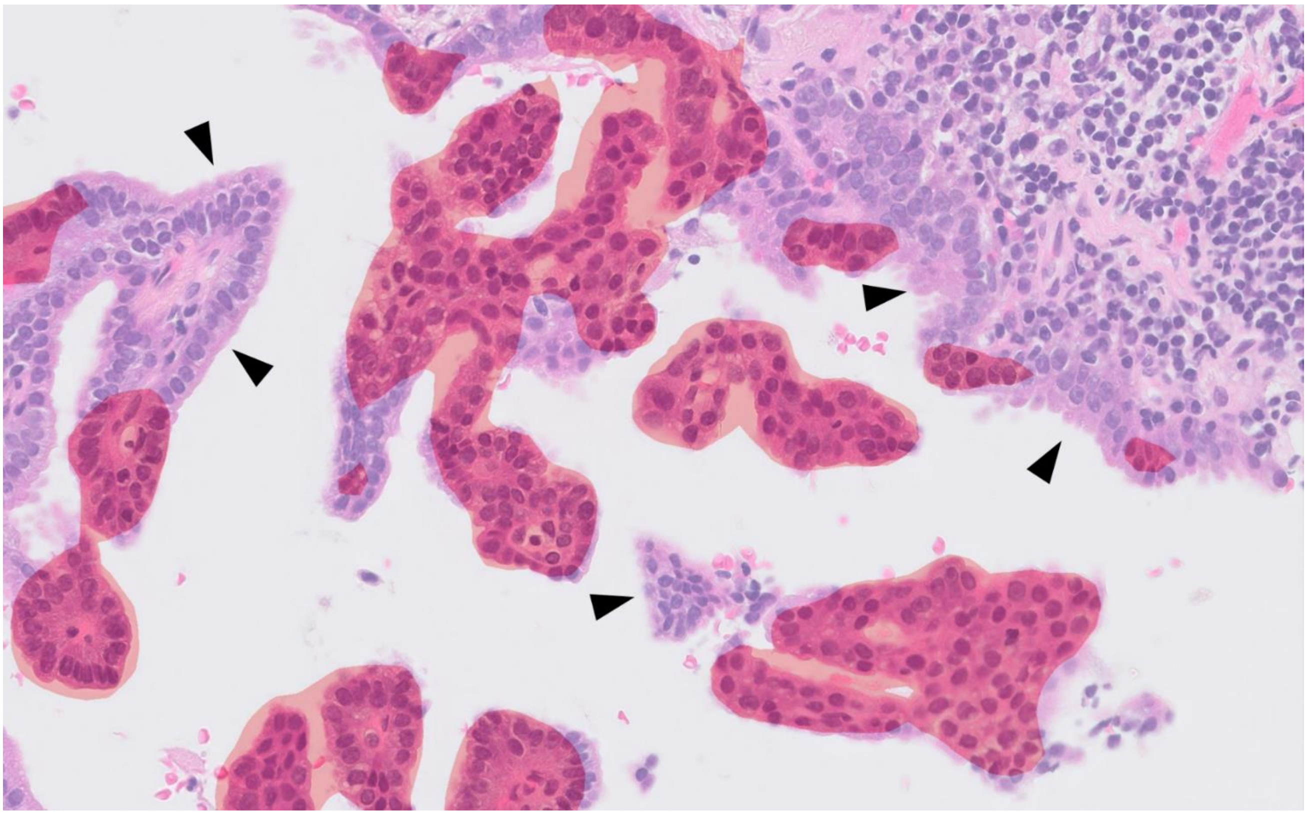

**Supplementary figure 2.** AI-generated carcinoma segmentation and pathologist adjustments.
Red masked regions were segmented and identified as carcinoma by AI. Certain areas were missed by AI (arrowhead). In such cases, the pathologists had to include the missing percentage to AI's calculation of tumor cellularity (AI-Score) to create the Adjusted-Score. In this ROI, the pathologists determined that approximately 30% had to be added to the AI-Score. Thus, 30% of the tumor area was missed by the segmentation algorithm. In practice, this decision was applied at the WSI level.

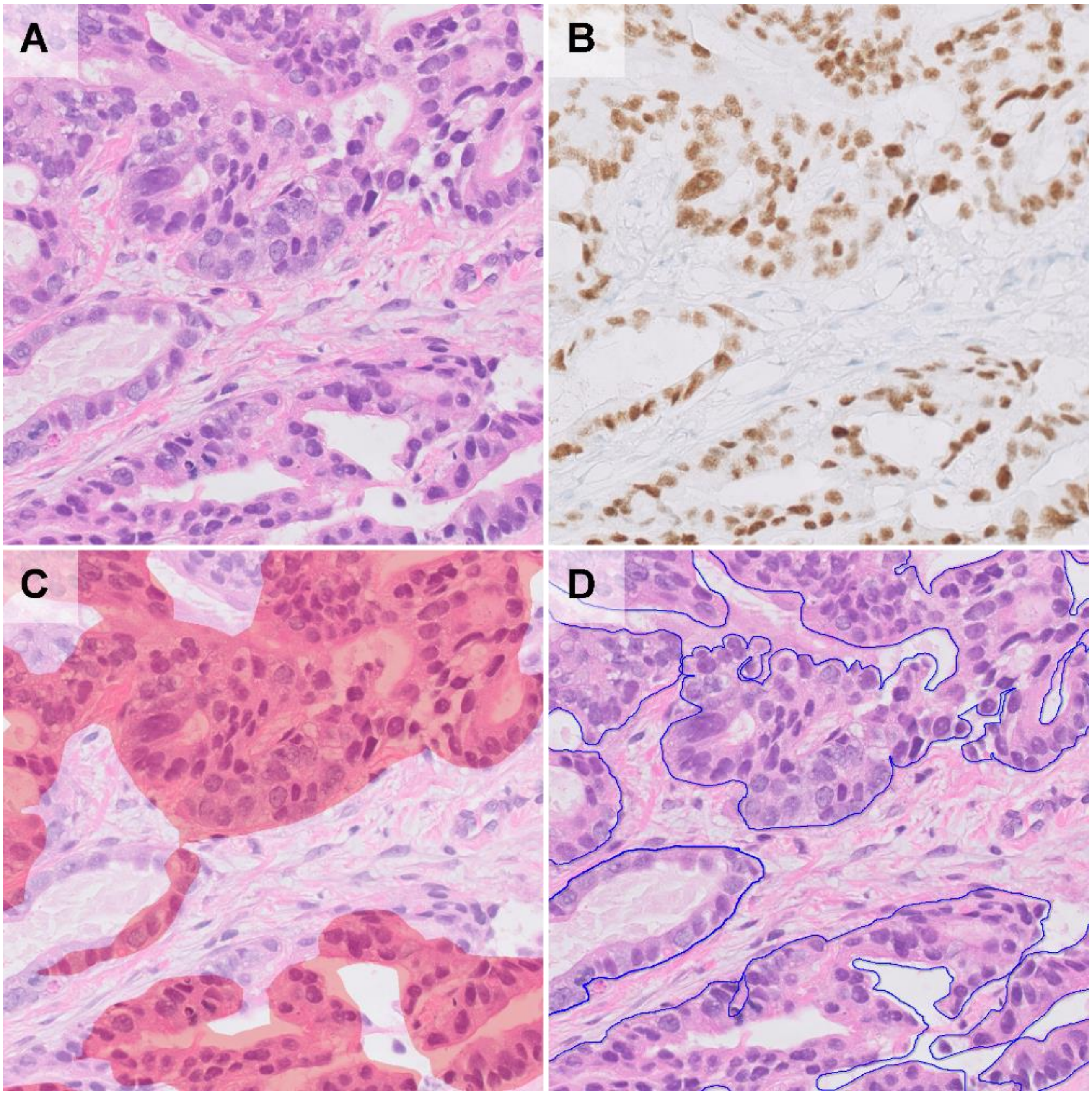


**Supplementary figure 3.** Creation of ground truth based on simultaneous observations of H&E slides and immunostained slides.

In 20 cases, pathologist-supervised annotations were collected to compile a ground truth dataset.

A) Original H&E image.

B) Re-stained slide of original H&E specimen with TTF-1 & napsin A cocktail.

C) Segmentation output by AI model.

D) Ground truth established by meticulous annotation with reference to immunostained slides. (40x).

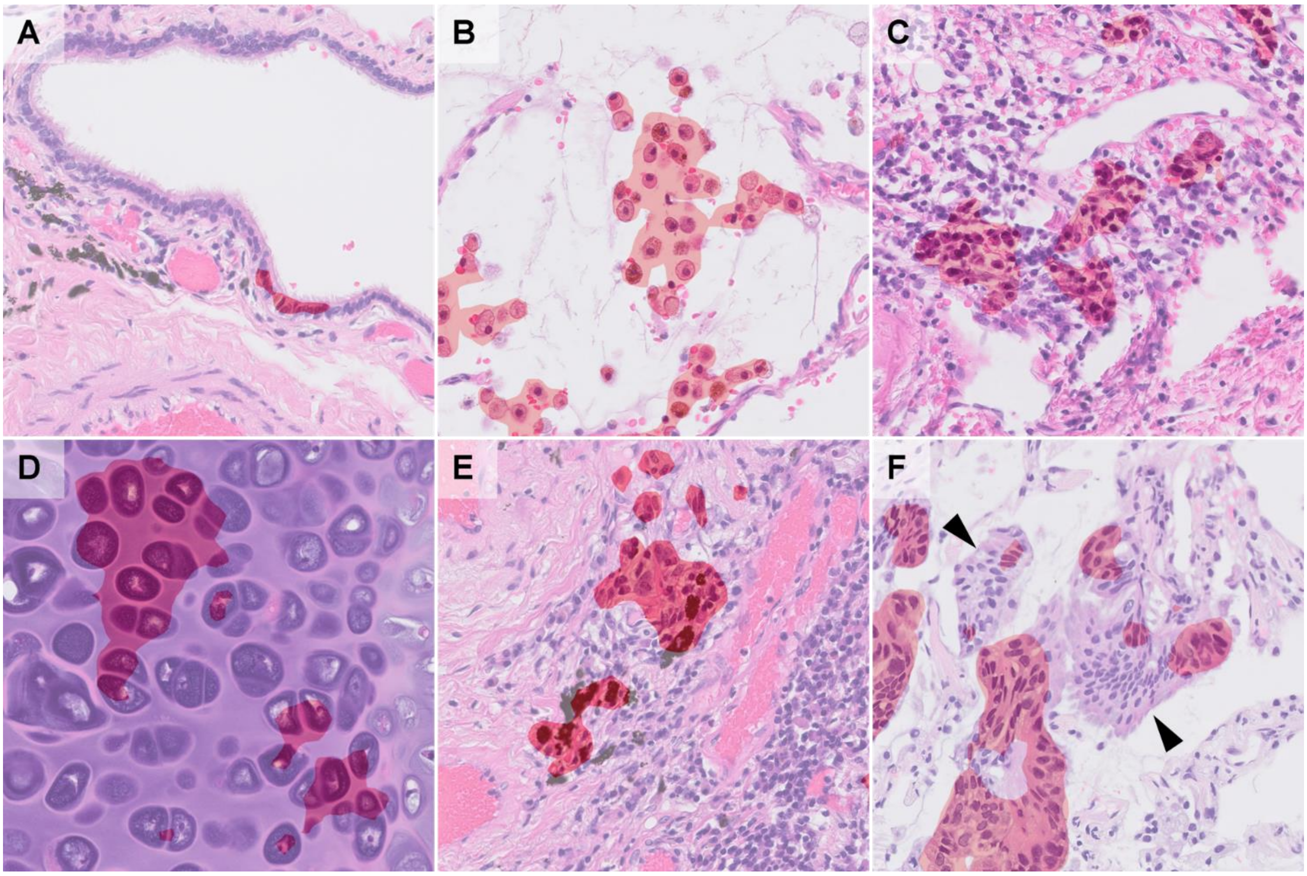


**Supplementary figure 4.** Examples of false positives and false negatives.

Bronchial epithelium (A), alveolar macrophages (B), lymphocyte infiltration/aggregation (C), tracheal cartilage (D), anthracotic pigments (E) were detected as false positives in certain instances. Invasive mucinous adenocarcinoma (F) tended to possess weakly atypical nuclei and could result in false negatives. Upon weighing the number of these false positives and false negatives, the pathologists considered the extent of correction actually required in AI-score. (40x)

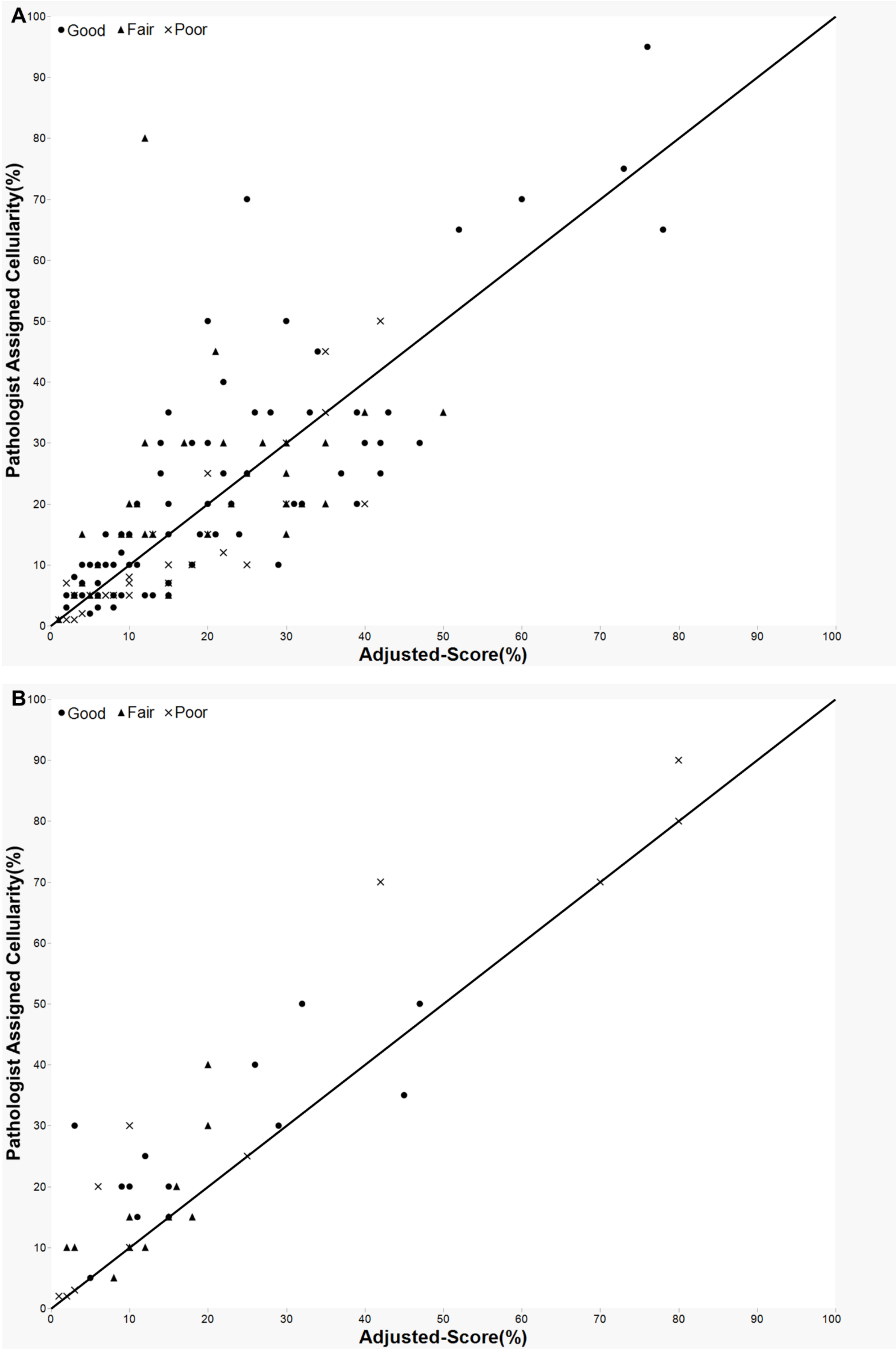

A
● Good ▲ Fair × Poor
Pathologist Assigned Cellularity(%)
Adjusted-Score(%)
0
10
20
30
40
50
60
70
80
90
100
B
● Good ▲ Fair × Poor
Pathologist Assigned Cellularity(%)
Adjusted-Score(%)

**Supplementary figure 5.** A tendency that was seen in the early and prospective phases.

These plots show the deviation between an individual pathologist's assigned scores (Path-Scores) and the final Adjusted-Score for 151 prospectively analyzed cases(A), and 37 retrospectively analyzed cases(B) in the early phase when AI was implemented into the cell count workflow. The line on both plots represents when Path-Score exactly equals the Adjusted-Score. Although the pathologist tended to overestimate the cellularity at first (points above the line), the frequency of overestimation and underestimation was almost even after using AI.

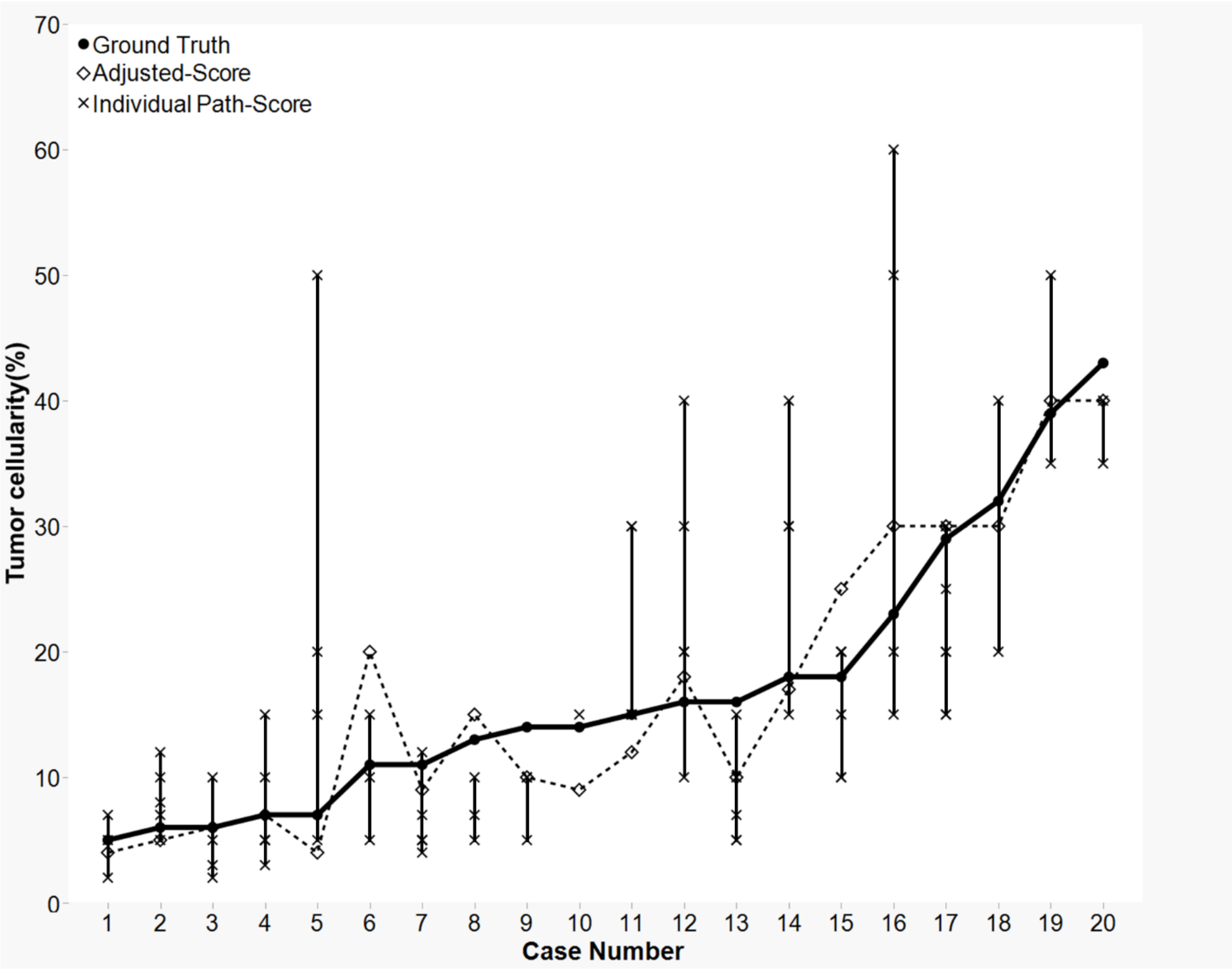


**Supplementary figure 6.** Deviation from Ground Truth in 20 cases.

Line plot showing Ground Truth (solid line), Adjusted-Score (dotted line), and Path-Scores (vertical line). As observed, the interobserver variability in the pathologists' estimations was resolved by adjusting AI-Score.

A

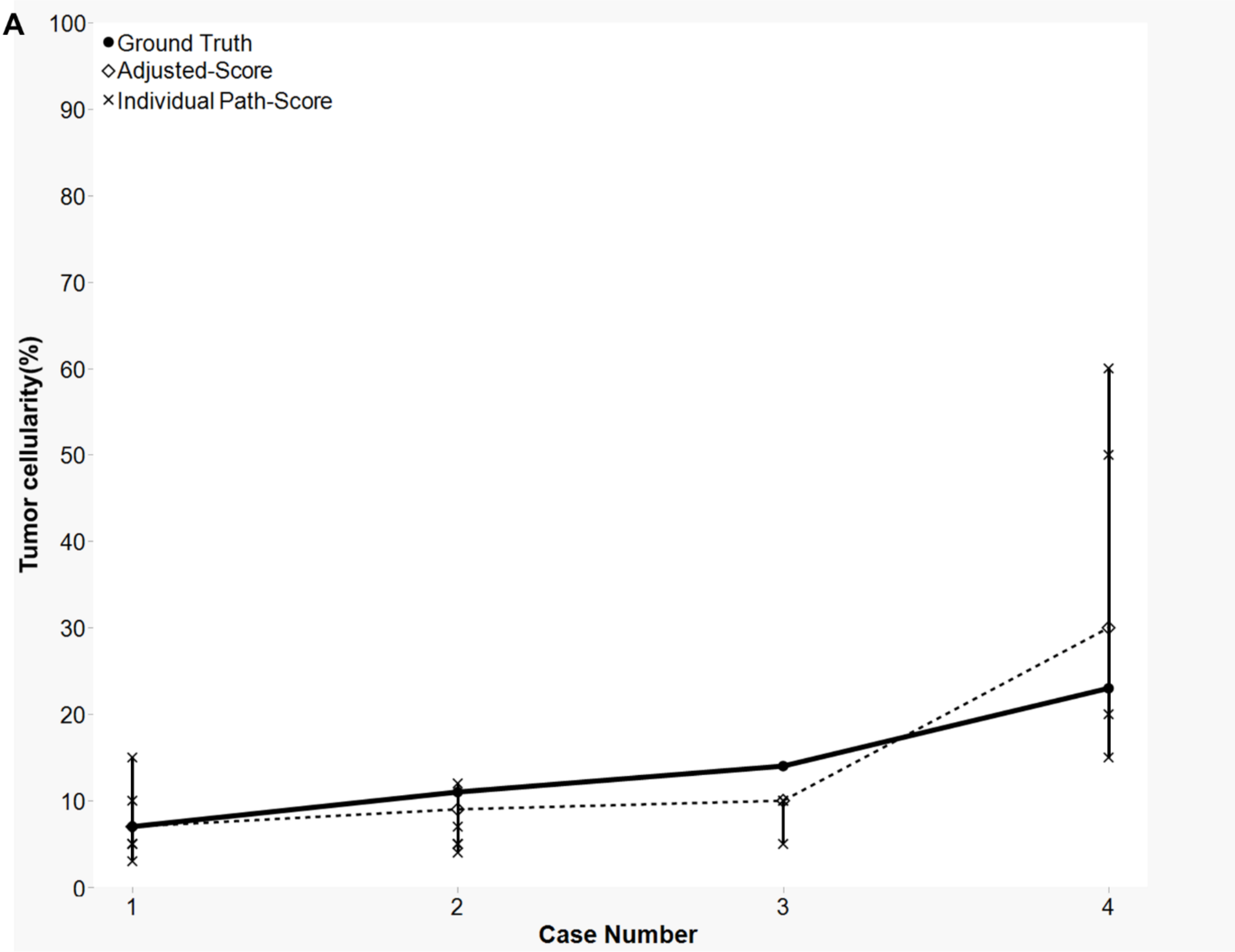

Ground Truth
Adjusted-Score
Individual Path-Score
Tumor cellularity(%)
100
90
80
70
60
50
40
30
20
10
0
1
2
3
4
Case Number

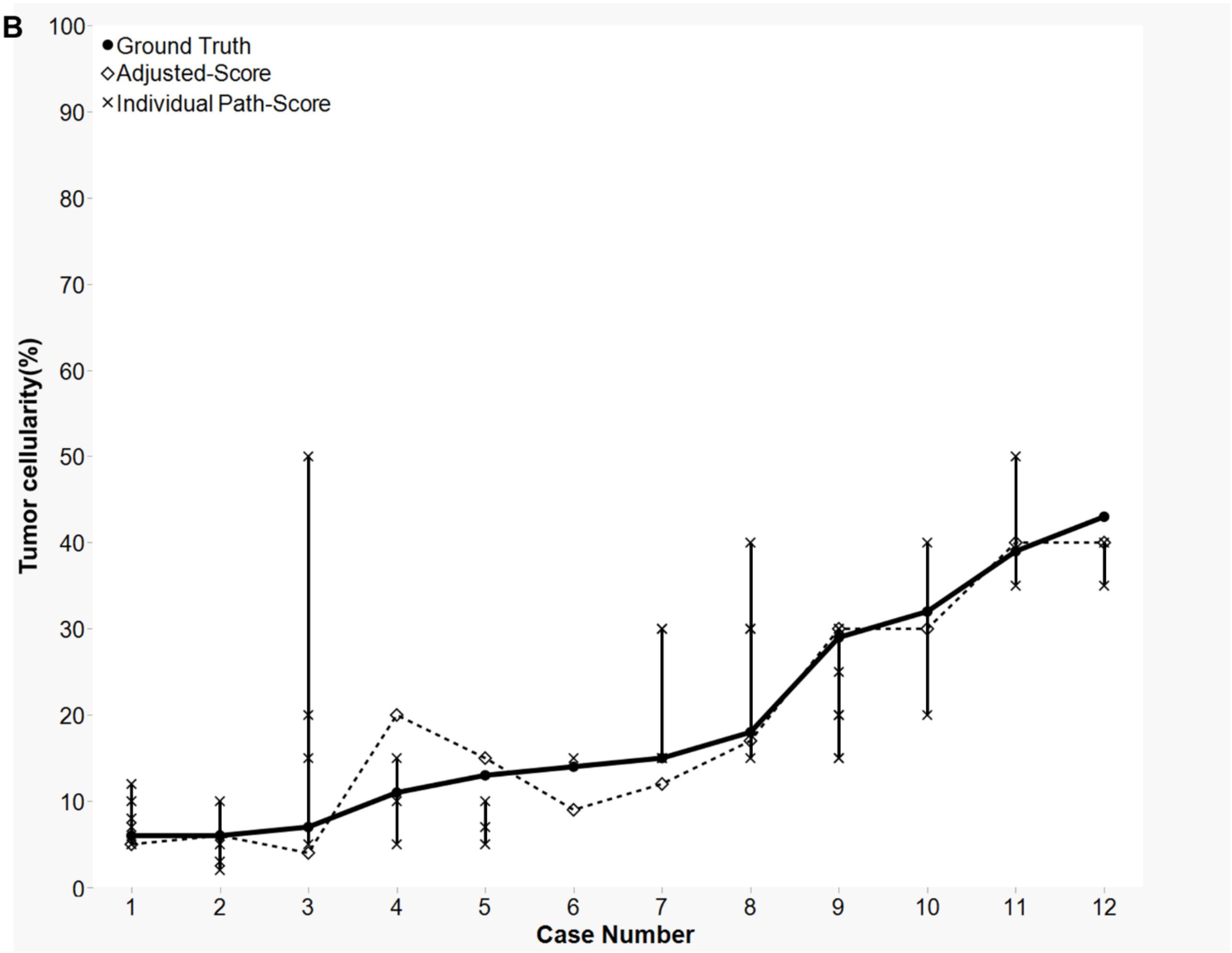
B
Ground Truth
Adjusted-Score
Individual Path-Score
Tumor cellularity(%)
0
10
20
30
40
50
60
70
80
90
100
1
2
3
4
5
6
7
8
9
10
11
12
Case Number

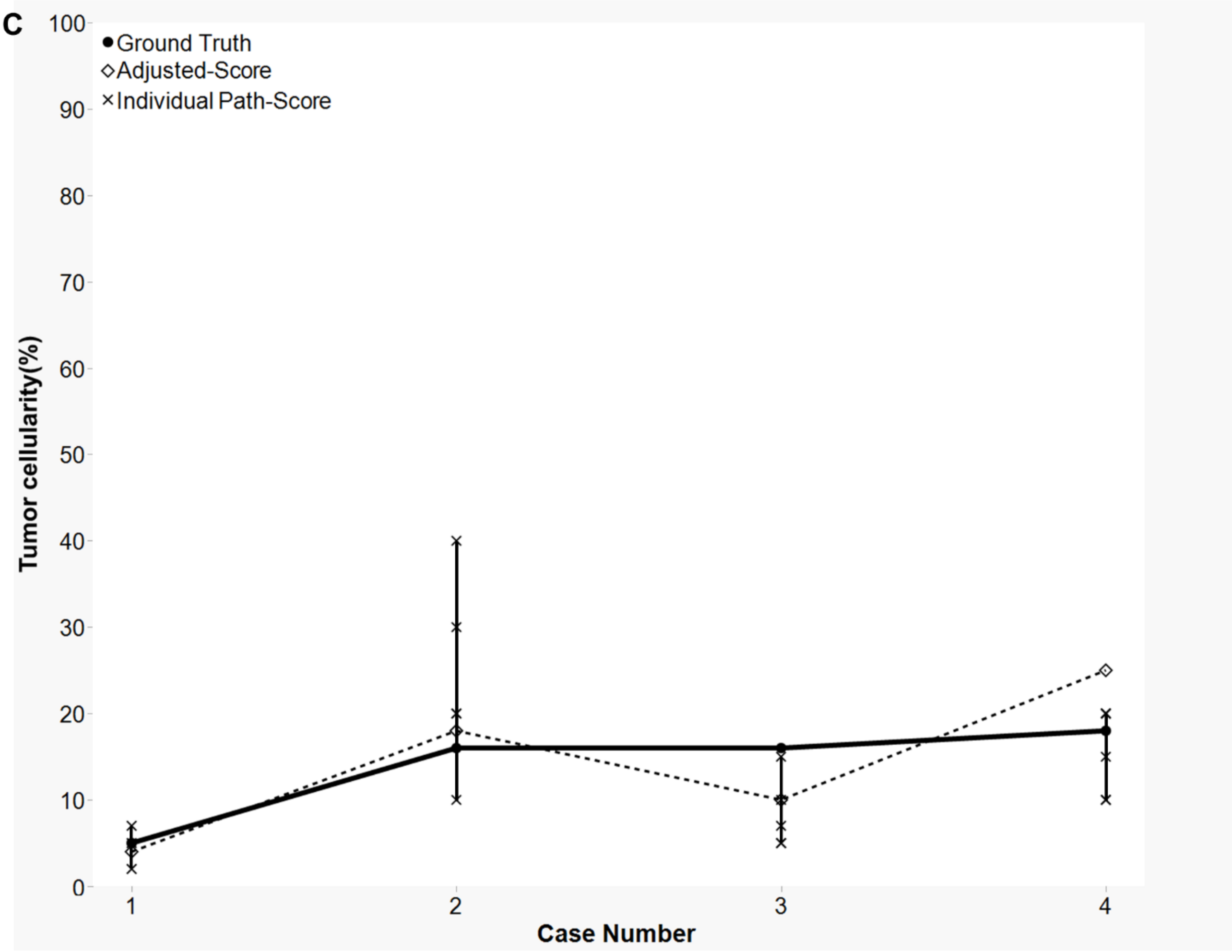


**Supplementary figure 7.** Deviation from ground truth per classification level.

Three line plots showing Ground truth (solid line), Adjusted-Score (dotted line), and individual Path-Score (vertical line) categorized as per the samples judged as good (a), fair (b), and poor (c).

**Supplementary Table 1.** Inter-observer variability of Path-Scores in the 29 cases judged as good in the retrospective phase

| No. | Ground Truth | AI-Score | Range of Path-Scores | Pathologist A | Pathologist B | Pathologist C | Pathologist D |
|---|---|---|---|---|---|---|---|
| 1 | 2 | 3 | 2–20 | 5 | 20 | 10 | 2 |
| 2 | 5 | 2 | 4–5 | 4 | 4 | 5 | 5 |
| 3 | 6 | 4 | 3–20 | 20 | 15 | 3 | 10 |
| 4 | 6 | 7 | 15–73 | 15 | 73 | 40 | 20 |
| 5 | 8 | 8 | 5–50 | 10 | 50 | 5 | 25 |
| 6 | 8 | 6 | 10–31 | 25 | 31 | 25 | 10 |
| 7 | 9 | 12 | 15–78 | 20 | 78 | 15 | 15 |
| 8 | 10 | 6 | 10–27 | 15 | 27 | 10 | 10 |
| 9 | 11 | 12 | 20–31 | 20 | 31 | 25 | 25 |
| 10 | 12 | 9 | 13–40 | 20 | 13 | 40 | 40 |
| 11 | 15 | 16 | 21–35 | 25 | 21 | 30 | 35 |
| 12 | 15 | 14 | 20–40 | 25 | 27 | 40 | 20 |
| 13 | 16 | 14 | 20–60 | 30 | 20 | 60 | 20 |
| 14 | 16 | 13 | 20–60 | 20 | 22 | 60 | 20 |
| 15 | 17 | 18 | 11–60 | 60 | 11 | 60 | 60 |
| 16 | 18 | 16 | 10–46 | 25 | 46 | 10 | 30 |
| 17 | 19 | 14 | 25–70 | 25 | 30 | 70 | 40 |
| 18 | 19 | 18 | 25–60 | 25 | 27 | 60 | 30 |
| 19 | 19 | 27 | 20–30 | 25 | 26 | 30 | 20 |
| 20 | 20 | 27 | 35–50 | 35 | 50 | 50 | 40 |
| 21 | 23 | 16 | 13–30 | 13 | 18 | 30 | 30 |
| 22 | 28 | 25 | 40–60 | 40 | 45 | 60 | 40 |
| 23 | 31 | 23 | 15–70 | 25 | 15 | 40 | 70 |
| 24 | 32 | 33 | 29–60 | 30 | 29 | 60 | 50 |
| 25 | 33 | 30 | 20–70 | 50 | 55 | 20 | 70 |
| 26 | 33 | 27 | 30–70 | 50 | 57 | 70 | 30 |
| 27 | 33 | 24 | 35–70 | 35 | 36 | 70 | 50 |
| 28 | 42 | 39 | 18–90 | 90 | 18 | 70 | 70 |
| 29 | 46 | 37 | 30–70 | 30 | 47 | 60 | 70 |

**Supplementary Table 2.** Mean adjustment from AI-Score per sampling method

| Sampling | Mean Adjustment | Standard Deviation |
|---|---|---|
| TBB/TBLB | 1.41 | 7.38 |
| CNB | 2.53 | 6.85 |
| Surgical | –0.29 | 9.96 |
| TBNA/TBAC | 2.33 | 4.15 |
| Cell Block | –19.33 | 30.90 |

TBB/TBLB: transbronchial biopsy/transbronchial lung biopsy; CNB: core needle biopsy; Surgical: surgical resection; TBNA/TBAC: transbronchial needle aspiration/transbronchial aspiration cytology

**Supplementary Table 3.** Classification level across three institutions

| Institute | N | Good | Fair | Poor |
|---|---|---|---|---|
| Nagasaki University Hospital | 26 | 73.1% (19) | 15.4% (4) | 11.5% (3) |
| Kameda Medical Center | 111 | 47.7% (53) | 27.9% (31) | 24.3% (27) |
| Awaji Medical Center | 14 | 57.1% (8) | 21.4% (3) | 21.4% (3) |
| Total | 151 | 53% (80) | 25% (38) | 22% (33) |
